# Early-Diverging SQR Enzyme in Antarctic Gloeobacterales Indicates Sulfide Tolerance in Thylakoid-Lacking Cyanobacteria

**DOI:** 10.1101/2025.10.24.684318

**Authors:** Louise Hambücken, Edi Sudianto, Jimmy H. Saw, Denis Baurain, Luc Cornet

## Abstract

Oxygenic photosynthesis, which converts solar energy into carbohydrates via a linear electron transport chain and two photosystems (PSII and PSI), first appeared in cyanobacteria approximately 3.3 Ga and drove the Great Oxidation Event around 2.4 Ga. During this period, euxinic conditions—characterized by sulfidic, anoxic oceans—posed a metabolic challenge to cyanobacteria, as sulfide inhibits PSII, the reaction center responsible for water splitting. Here, we report the presence of an early-diverging form of the sulfide quinone reductase (SQR) enzyme in Antarctic representatives of *Gloeobacterales*, the earliest-branching cyanobacterial lineage lacking thylakoids. Phylogenetic analyses reveal that these SQR sequences are the earliest-diverging cyanobacterial SQR known to date, predating the multiple lateral gene transfer events previously observed in the phylum. Additional searches in metagenomic datasets indicate that such sequences are restricted to cold environments. Our findings unveil possible adaptive strategies of early cyanobacteria to cope with sulfidic stress and point to Antarctic lakes as preserved natural laboratories for investigating cyanobacterial diversification and the evolution of oxygenic photosynthesis under euxinic conditions.

**Impact Statement:** The diversification of cyanobacteria during and after the Great Oxidation Event occurred in early Proterozoic oceans that were partially euxinic (anoxic and sulfidic) a condition generally considered incompatible with oxygenic photosynthesis due to photosystem II inhibition. The presence of a sulfide quinone reductase in an Antarctic early diverging cyanobacterium lacking thylakoids gives credit on an ancestral evolutionary stage where oxygenic and anoxygenic traits coexisted within cyanobacteria. The occurrence of these organisms in Antarctic lakes under euxinic conditions offers a natural laboratory for studying the physiology and adaptation of the first oxygenic photosynthetic organisms.

## Main

Oxygenic photosynthesis is a biochemical process that converts solar energy into carbohydrates by transferring electrons from water through a linear electron transport (LET) chain involving two photosystems (PSII and PSI). This process first appeared in cyanobacteria approximately 3.3 Ga (1), releasing oxygen into both the oceans and the atmosphere as a byproduct, and ultimately leading to the Great Oxidation Event (GOE) around 2.4 billion years ago (2). When oxygen accumulated in the atmosphere, it created oxidizing conditions that promoted the weathering of terrestrial sulfides, as evidenced by the presence of evaporites (sulfate-rich rocks) dated between 2.7 and 2.2 Ga (3). This weathering released large quantities of sulfate into the oceans, which were then used by sulfate-reducing bacteria, producing sulfide in the process (4). The accumulation of sulfide transformed the oceans into an anoxic and sulfidic environment—known as the euxinic ocean—as proposed by Canfield (1998) (5). The Fe^2^□ present in the oceans, which had first reacted with oxygen to form ferric hydroxides and banded iron formations characteristic of oceanic oxygenation, began instead to bind with sulfur, forming pyrite (FeS) (4). According to Canfield (1998), these euxinic conditions were widespread during early and mid-Proterozoic (5), though subsequent studies tempered this view (4). Johnston et al. (2006) nevertheless demonstrated that H□S was sufficiently abundant in the oceans for the chemocline to reach the surface, releasing H□S into the atmosphere and sometimes impacting the entire water column at the end of the first era of the Proterozoic, the Paleoproterozoic (6). As a result, cyanobacteria living in the photic zone of the mid-Proterozoic experienced alternating oxygenated and sulfidic conditions (7). The existence of euxinic environments during cyanobacterial diversification in the Proterozoic likely posed a significant challenge for these organisms, since sulfide inhibits photosystem II by an as-yet-unclear mechanism (8, 9).

In most cyanobacteria, oxygenic photosynthesis takes place in specialized membrane compartments known as thylakoids. However, the earliest-diverging cyanobacterial order, *Gloeobacterales*, lacks thylakoids, and its LET chain is instead located in the cytoplasmic membrane, where oxygenic photosynthesis occurs (10). Given the ancestral nature of *Gloeobacterales*, the emergence of oxygenic photosynthesis must have taken place within the cytoplasmic membrane before the relocation of the LET chain to the thylakoids (11). *Gloeobacterales* have been identified in a variety of environments, including cold, wet-rock and low-light environments (12). Within their known diversity, there exists a group of taxa represented exclusively by metagenome-assembled genomes (MAGs) that are endemic to Antarctica.

During the Proterozoic, anoxygenic photosynthesis—which, according to Nishihara *et al*. (2024) (13), is thought to have preceded oxygenic photosynthesis—was widespread. This type of photosynthesis notably involves the enzyme sulfide-quinone reductase (SQR), which catalyzes H□S oxidation, supplying electrons to the LET chain. Alongside the purple sulfur bacteria (e.g., *Chromatium*) and green sulfur bacteria (e.g., *Chlorobi*), for which sulfide oxidation is the sole photosynthetic pathway, some cyanobacteria are also capable of using the SQR enzyme intermittently (14, 15). In this study, we report the presence of an early diverging form of the SQR enzyme in Antarctic representatives of the *Gloeobacterales*.

We built a profile of *Gloeobacterales*-specific SQR Type I (SQRI) using the protein sequences derived from three Antarctic Gloeobacterales MAGs (GCA_949127895.1, GCA_949127685.1, GCA_038245785.1) and performed an orthology search against 107,237 representative genomes from the GTDB database (16) (**Supplementary Figure 1**). This search yielded 19,081 homologous sequences, out of which we computed a phylogenetic tree **(Supplementary Figure 2**) where cyanobacteria form a monophyletic group composed exclusively of the SQRI gene. Interestingly, the three *Gloeobacterales* SQRI from Antarctica represent the earliest-diverging SQRI sequences among this cyanobacterial subgroup, consistent with the phylogenetic position of *Gloeobacterales* within the cyanobacterial lineage. Previous studies have reported lateral gene transfers (LGTs) among cyanobacteria, notably involving *Thermostichales* (17), the second cyanobacterial order to diverge. In a smaller tree inferred from 192 cyanobacterial sequences and rooted on *Gloeobacterales* SQRI (**Figure 1**), several well recognized cyanobacterial orders of Strunecký *et al*. (2023) (18) (e.g., Oscillatoriales, Nostocales) are exploded and intermingled with strong bootstrap support, indeed evidencing pervasive LGT in the SQR evolution. Nevertheless, the three *Gloeobacterales* sequences represent an early-diverging version of the cyanobacterial SQRI, rather than a secondary acquisition, since extensive orthology searches (**Supplementary Figure 1**) only returned a single closely related sequence. This sequence originates from a *Rhizonema* MAG (GCA_029379215.1) recovered from a cyanolichen inhabiting a cold alpine environment, at 1,600 m elevation (personal communication, Botanical Research Institute of Texas).

**Figure 1.**
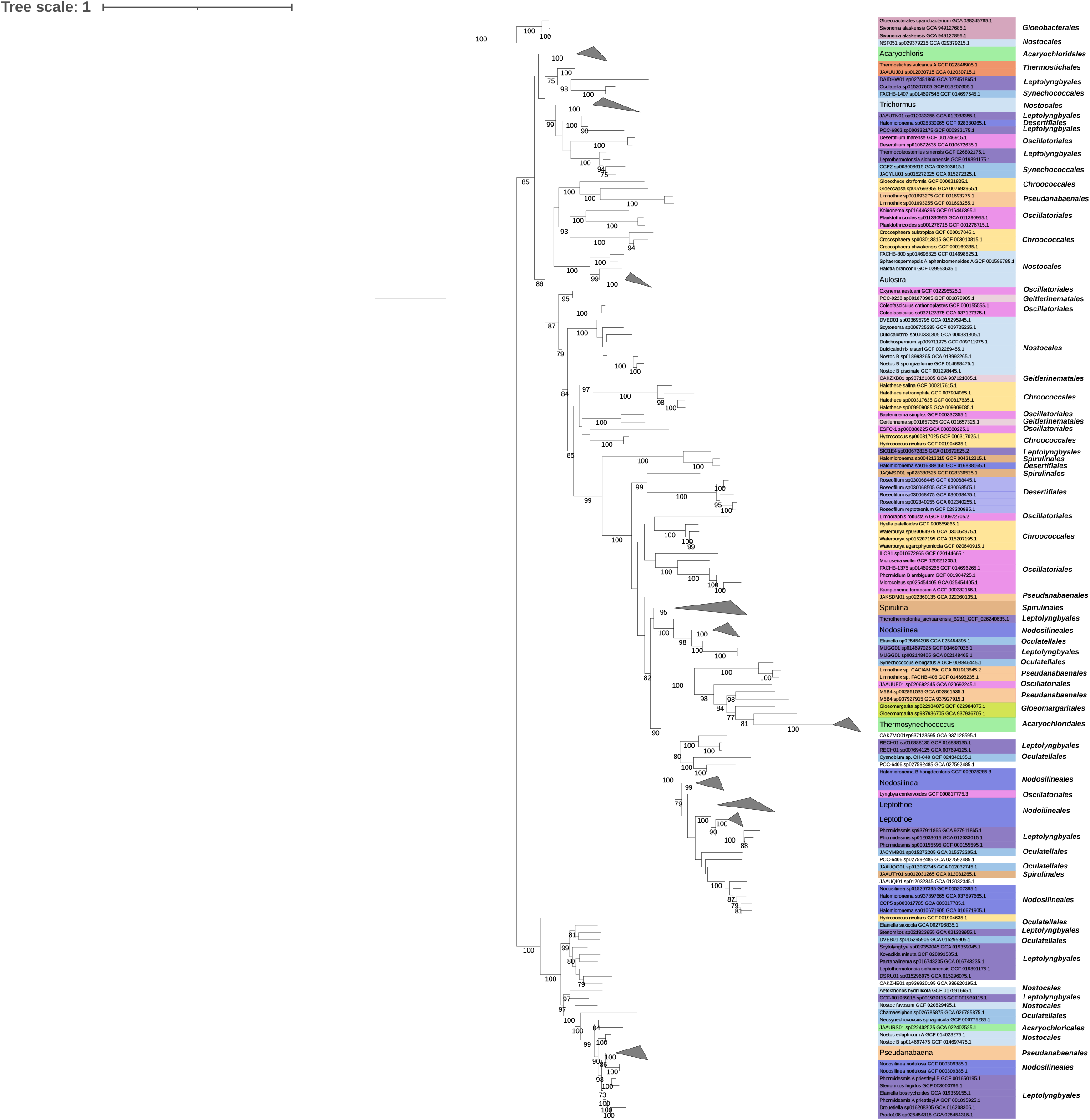
Phylogenetic tree of Cyanobacteria SQRI sequences. The tree, constructed from 192 sequences across 438 unambiguously aligned positions, was inferred using IQ-TREE with the best-fit model (Q.PFAM+I+R5). It is rooted on the *<u>Gloeobacterales</u>* sequences. The classification proposed by Strunecký et al. (2023) was applied to the genomes, using the same color code as in that study. The first appearance of a Strunecký et al. (2023) order is indicated on the tree. Only bootstrap proportions above 70% are shown.

**Figure 2.**
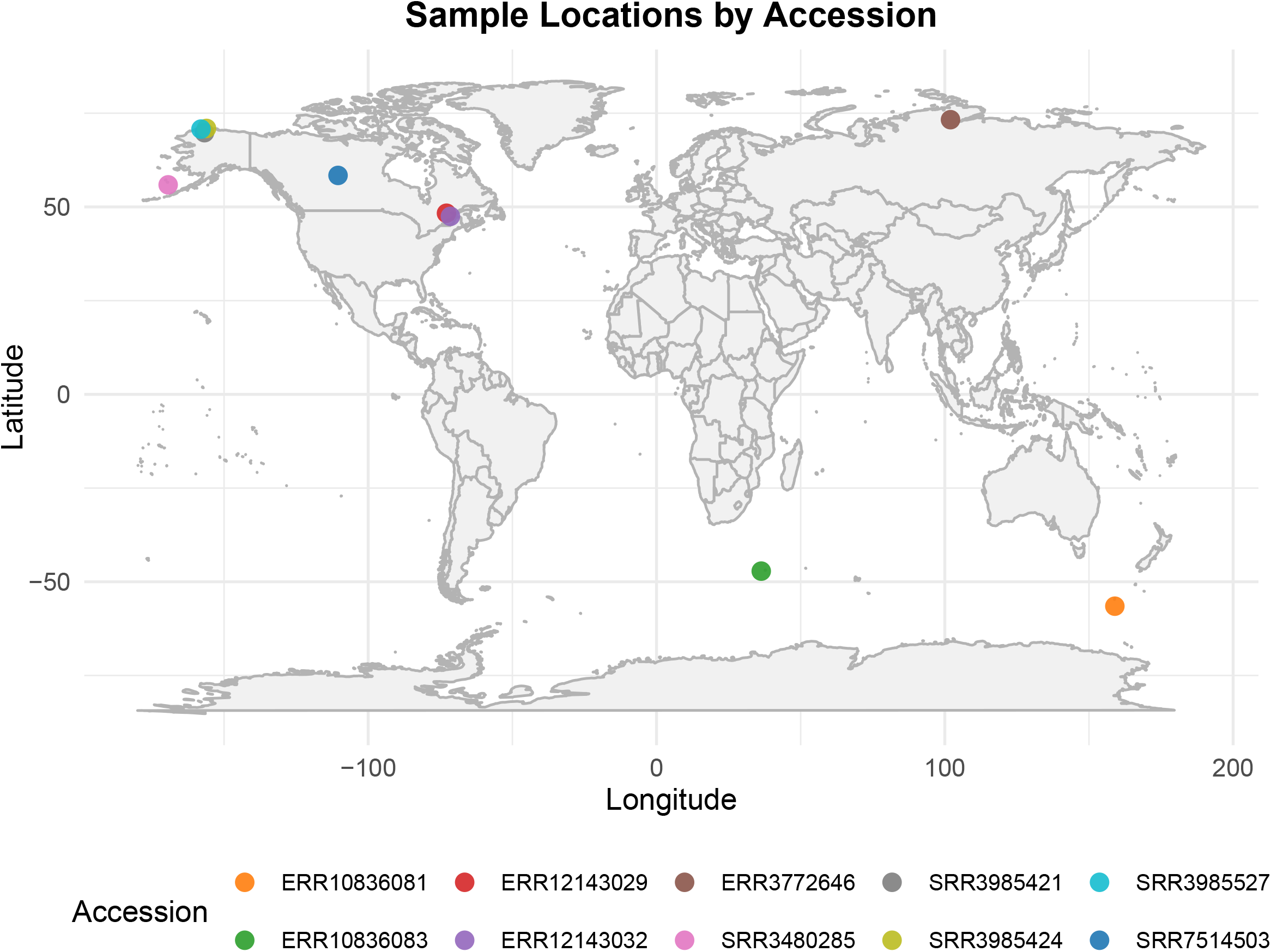
Geographic localization of the SRA datasets identified by Logan.

A BLASTP search using *Gloeobacterales* SQRI sequences against the NCBI GenBank database yielded no additional results. Using Logan (19), we performed a nucleotide kmer based search using SQRI sequences from *Gloeobacterales* as queries to detect kmers specific to the *Gloeobacterales*-type SQRI across the largest DNA sequencing dataset, the sequence read archive of the National center for Biotechnology information (NCBI SRA). This analysis returned 11 matches, mostly (9/11) originating from cold environments (subarctic, subantarctic, and polar) (**Figure 2; Supplementary Table 1**). The SQRI sequences from these SRAs could not be fully assembled into metagenomes or binned into MAGs; only partial hits were recovered. Nevertheless, the specificity of the kmers used by Logan (19) ensures the presence of these sequences within the SRAs. The SQRI sequence of Antarctic *Gloeobacterales* MAGs therefore appears to be the earliest-diverging cyanobacterial SQRI sequence reported to date, having diverged from the ancestral sequence to adapt to cold environments. In many cases, Antarctic lakes are meromictic and exhibit strong water column stratification, notably due to prolonged ice cover (e.g., (20)). The resulting limited water mixing leads to the accumulation of sulfide, thereby exposing the microbial mats of these lakes to euxinic conditions. Although the precise conditions in these lakes differ from those of the Proterozoic oceans, the presence of cyanobacteria carrying an ancestral version of the LET chain alongside an early-diverging SQRI suggests that Antarctic lakes are a valuable natural system for studying cyanobacterial diversification under Proterozoic conditions.

Recovery of relatively few metagenomic sequence reads of *Gloeobacterales* in the Antarctic lake samples indicates that they are components of the “rare biosphere”, hypothesized to be highly diverse but only present in low abundance (21, 22). This makes it challenging to study them using either sequence-or cultivation-based approaches as their genomes cannot be easily constructed unless ultra-deep sequencing is applied or they outgrow other fast-growing cyanobacteria in cultivation experiments—for example, species of *Gloeobacter* are notoriously slow-growing (23, 24). Yet, despite several billion years of evolution, they have not gone extinct and remain integral, albeit low-abundance, components of various ecosystems, potentially serving as sources of genetic diversity in Antarctic lake biofilm communities.

The SQRI sequences reported in this study possess both the sulfur-binding sites and the FAD/NADP-binding domains, catalyzing sulfur transfer and reduction processes using FAD/NADP as cofactors. (**Supplementary Figure 3**). Since *Gloeobacterales* lack thylakoids, their periplasmic space is considered homologous to the thylakoid lumen (25). Consequently, they very likely are the only SQR enzymes active within the periplasmic space of cyanobacteria, thus mimicking the ancestral cyanobacterial LET chain subjected to the euxinic conditions of the Proterozoic. Indeed, the presence of an SQR must allow these organisms to continue generating energy via photosynthesis in the absence of a functional PSII, as the earliest cyanobacteria likely did.

## Contributions

LH performed all analyses at the exception of Logan search. ES constructed the GTDB dataset and performed the Logan search. LH, ES, JS, DB, and LC wrote the manuscript. All authors read and approved the final version of the manuscript.

## Acknowledgments

We thank Manuel Dal Forno from the Botanical Research Institute of Texas for providing information regarding the *Rhizonema* genome. This work was supported by a research grant (PDR T.0018.24 OR-OX-PHOT-IN-CYN) from the Belgian National Fund for Scientific Research (F.R.S.-FNRS) to DB. LC is supported by a mandate from the Belgian National Fund for Scientific Research (F.R.S.-FNRS).

