## Supplementary material for "Early-Diverging SQR Enzyme in Antarctic Gloeobacterales Indicates Sulfide Tolerance in Thylakoid-Lacking Cyanobacteria": Supp file 1

**Materials and methods**

**Inference of SQR type I orthologous sequences**

We used OrthoFinder v2.5.5 (1) to identify the orthologous group corresponding to the SQRI sequence. To work around OrthoFinder’s memory limit of 950 genomes, we first performed a dereplication of the 45316 representative genomes of Terrabacteria species downloaded from GTDB r220 (2). The purpose of the dereplication was to select the most complete and least contaminated genomes while preserving the diversity of Terrabacteria in GTDB.

Genomes that passed the GUNC (3) contamination test were grouped by GTDB taxonomy order and ranked based on their CheckM2 (4) completeness minus CheckM2 contamination score. If an order contained only one genus, the two best genomes were selected; otherwise, one genome was chosen per order. This resulted in 1452 genomes without Cyanobacteria, which were added separately. The genome set for the Cyanobacteria phylum was sampled more extensively, resulting in 432 genomes. The list of cyanobacterial genomes was obtained from GTDB release r220 (2). One to two representative genomes were selected for each genus, with preference given to assemblies exhibiting high completeness (>80%) and low contamination (<10%) as assessed by CheckM2 (4). To ensure adequate representation of basal cyanobacterial lineages, all available genomes from NCBI corresponding to the orders *Gloeobacterales*, *Thermostichales*, *Pseudanabaenales*, and *Gloeomargaritales* were included. The *Pseudanabaenales* dataset was subsequently dereplicated using ToRQuEMaDA (version 1) (5). The proteomes of the 1884 genomes were obtained from the NCBI database when available and otherwise predicted using Prodigal v2.6.3 (6). A ribosomal tree was subsequently inferred using the GEN-ERA toolbox (7), using IQ-TREE v2.4.0 (8) with the best model parameter and 1000 ultrafast bootstrap replicates. The ribosomal tree was then pruned using PARNAS v0.1.6 (9) by selecting 950 representative species, covering 90% of the overall diversity based on the generated diversity scores.

The orthogroup, computed with OrthoFinder v2.5.5 (1) on the 950 representative species, corresponding to SQR type I was identified based on sequence similarity with the *Synechocystis* sp. PCC 6803 substr. Kazusa Sll5036 (NCBI Protein Accession Number: BAD01806.1) using BLASTp searches (BLAST v2.9.0+ (10)). A phylogenetic tree of the SQR orthogroup was then generated using IQ-TREE v3.0.0 (8) with the best-fit model parameter (LG+I+G) and 1000 ultrafast bootstrap replicates. The sequences were aligned using MAFFT v7.505 (11). The sites of the alignment were conserved if they had at most 30% gaps and the sequences with a minimum of 30% length were exported using ali2phylip.pl (D. Baurain; <https://metacpan.org/release/Bio-MUST-Core>). To avoid problems of paralogy during orthologous sequence inference, the sequences corresponding to SQR type I were located within the gene tree based on their sequence similarity to the reference sequence Sll5036. A profile HMM (HMMER v3.4 (12)) was then generated from 11 cyanobacterial SQR type I orthologous sequences. This HMM profile was then used to search the proteomes of 432 Cyanobacteriota genomes in order to increase the sampling beyond the 950-sequence limit imposed by OrthoFinder’s memory constraints. This resulted in the discovery of three SQR type I sequences in *Gloeobacter* MAGs.

**SQR phylogenetic tree inference using GTDB representative species**

An HMM search was conducted across a dataset containing the proteomes of all representative species (107237) from GTDB release r220 (2) and the proteomes of the three Gloeobacter MAGs. The proteomes of the selected genomes were inferred using Prodigal v2.6.3 (6). A preliminary tree was generated based on the sequences selected according to the criteria listed in **Supplementary Table 1** with the Ompa-Pa tool v0.251830 (D. Baurain; <https://metacpan.org/dist/Bio-MUST-Apps-OmpaPa>) using the default parameters of the protein mode of FastME v2.1.6.4 (13) and FastTree v2.1.11 (14). The selection was visualized with the Ompa-Pa tool based on taxonomy, alignment coverage and number of copies per genome (**Supplementary Figure 1**). This tree (**Supplementary Figure 2**) contains a subtree represented exclusively by cyanobacterial SQRI sequences. A phylogenetic tree of the proteins contained in this subtree corresponding was subsequently performed using IQ-TREE (8) with best-fit model (Q.PFAM+I+R8) and 1000 ultrafast bootstrap replicates (**Figure 1**). For each of the trees, the sequences were aligned using MAFFT v7.505 (11). The sites of the alignment were conserved if they had at most 30% gaps and the sequences with a minimum of 30% length were exported using ali2phylip.pl (D. Baurain; <https://metacpan.org/release/Bio-MUST-Core>). The sequence similarity between the reference SQR type I (Sll5036) and the selected proteins was assessed with a BLASTp search (10).

**Contamination assessment of the three *Gloeobacter* MAGs**

The three Gloeobacterales MAGs were subjected to contamination assessment using GUNC (3) and CheckM2 (4) within the GEN-ERA toolbox, and the results are summarized in **Supplemental Tables 3 and 4**.

**Functional predictions of Gloeobacterales SQR type I**

Functional domains were predicted using the web-based version of InterProScan 5 v106.0 (15). The multiple sequence alignment in Supplemental Figure 4 was annotated based on 10.1021/bi9003827.

**Logan kmer search**

Logan SRA search was performed using the first 1,000 bp of both Candidatus *Sivonenia alaskensis* GCA_949127895.1 and GCA_949127685.1 as queries in Logan Search (https://logan-search.org/; (16)). The search was performed against the “All” database using the default similarity threshold (0.5).

**Supplemental Figures and Tables**

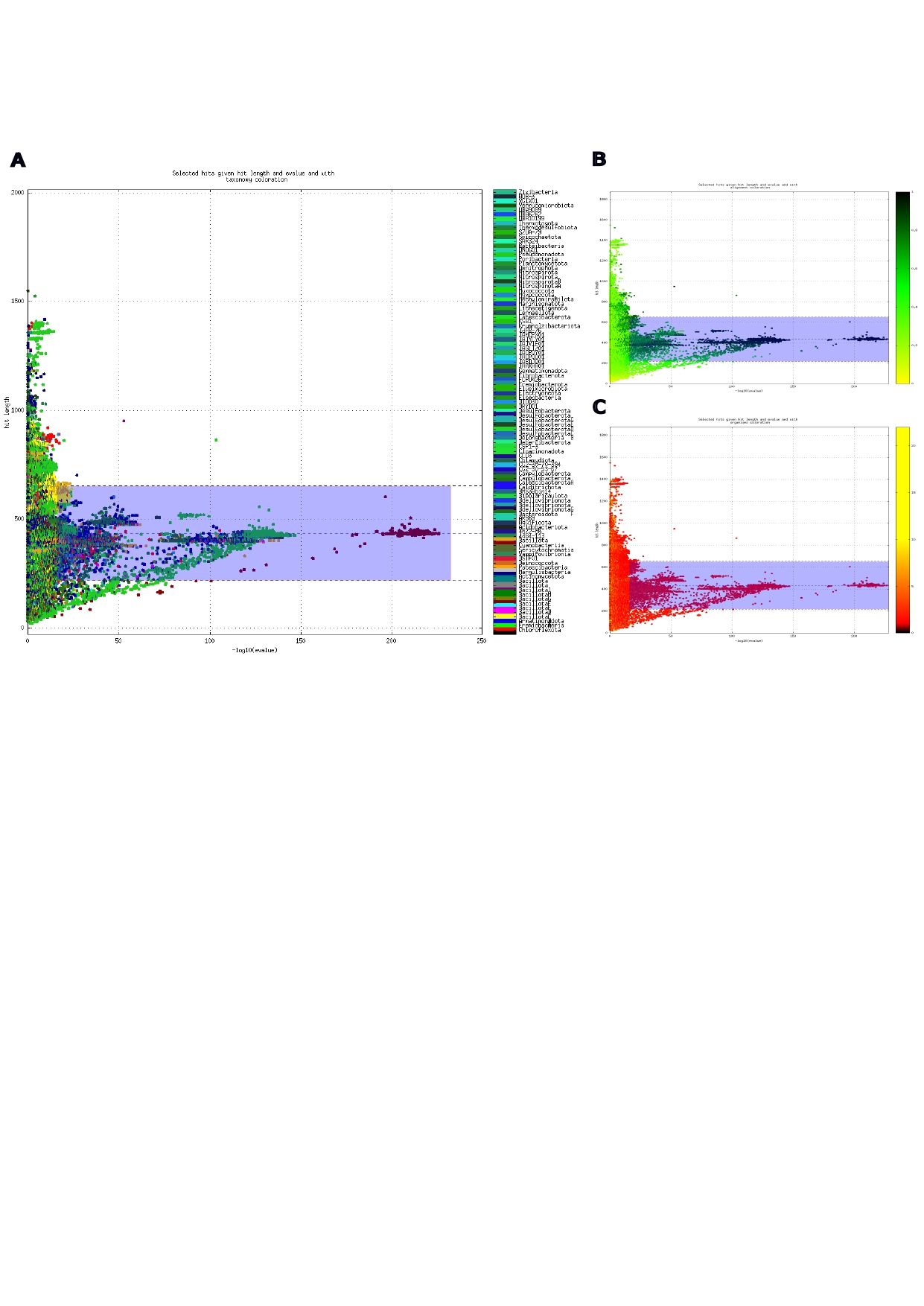

**Supplemental Figure 1: Visualization of the selection of the sequences similar to Gloeobacterales SQR type I.** Matching sequences were plotted based on their sequence length in function of their -log(e-value) score. The blue box corresponds to the selection. The sequences were visualized based on their (A) taxonomy, (B) alignment coverage and (C) number of copies per genome.

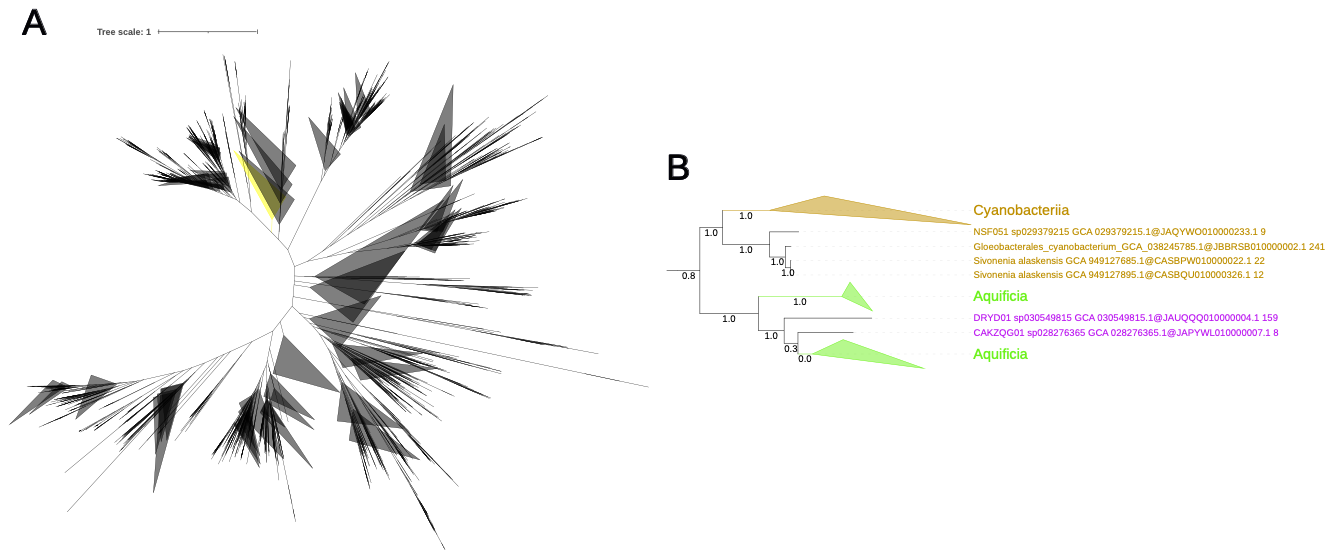

**Supplemental Figure 2: SQR phylogenetic tree prior to selection.** (A) Preliminary tree. (B) Subtree containing Cyanobacterial SQR type I sequences. The tree was constructed from 19081 sequences, corresponding to 453 unambiguously aligned positions, using  FastTree with default parameters. The subtree highlighted in yellow represents the selected SQR type I subtree, corresponding to 192 sequences. The local support values are based on the Shimodaira-Hasegawa (SH) test ([10.1093/oxfordjournals.molbev.a025808](https://doi.org/10.1093/oxfordjournals.molbev.a025808)).

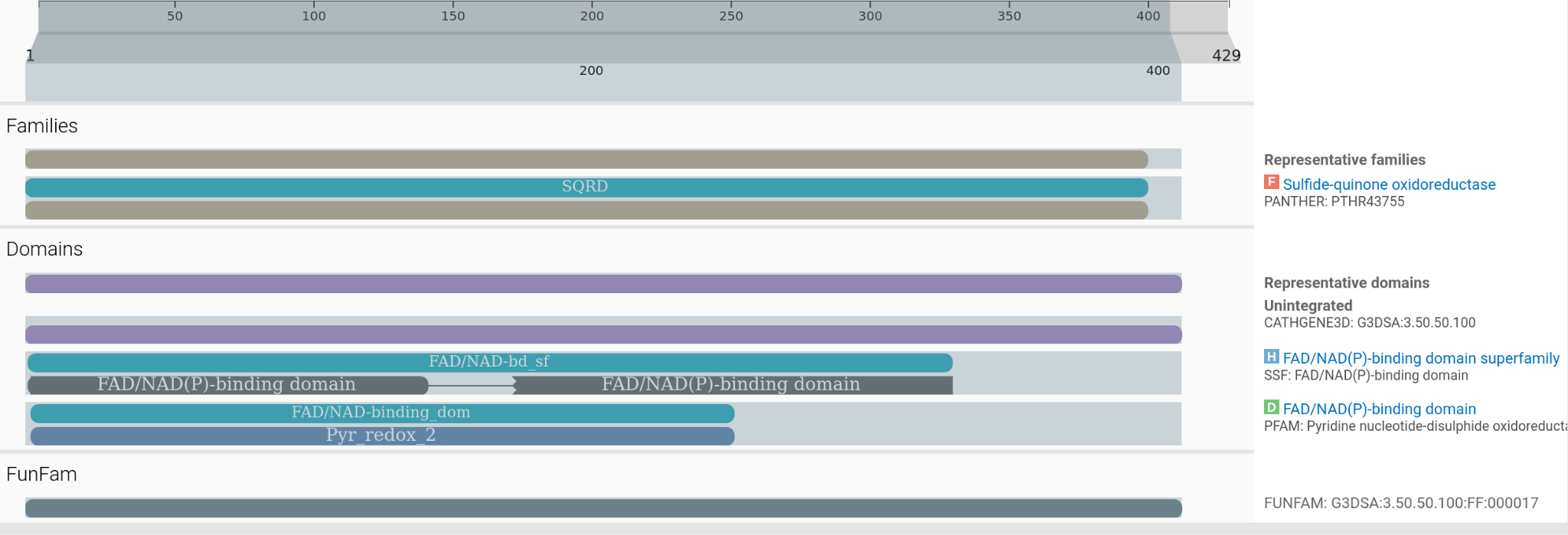

**Supplemental Figure 3: Functional domains of the SQRI sequence from GCA_949127895.1.**

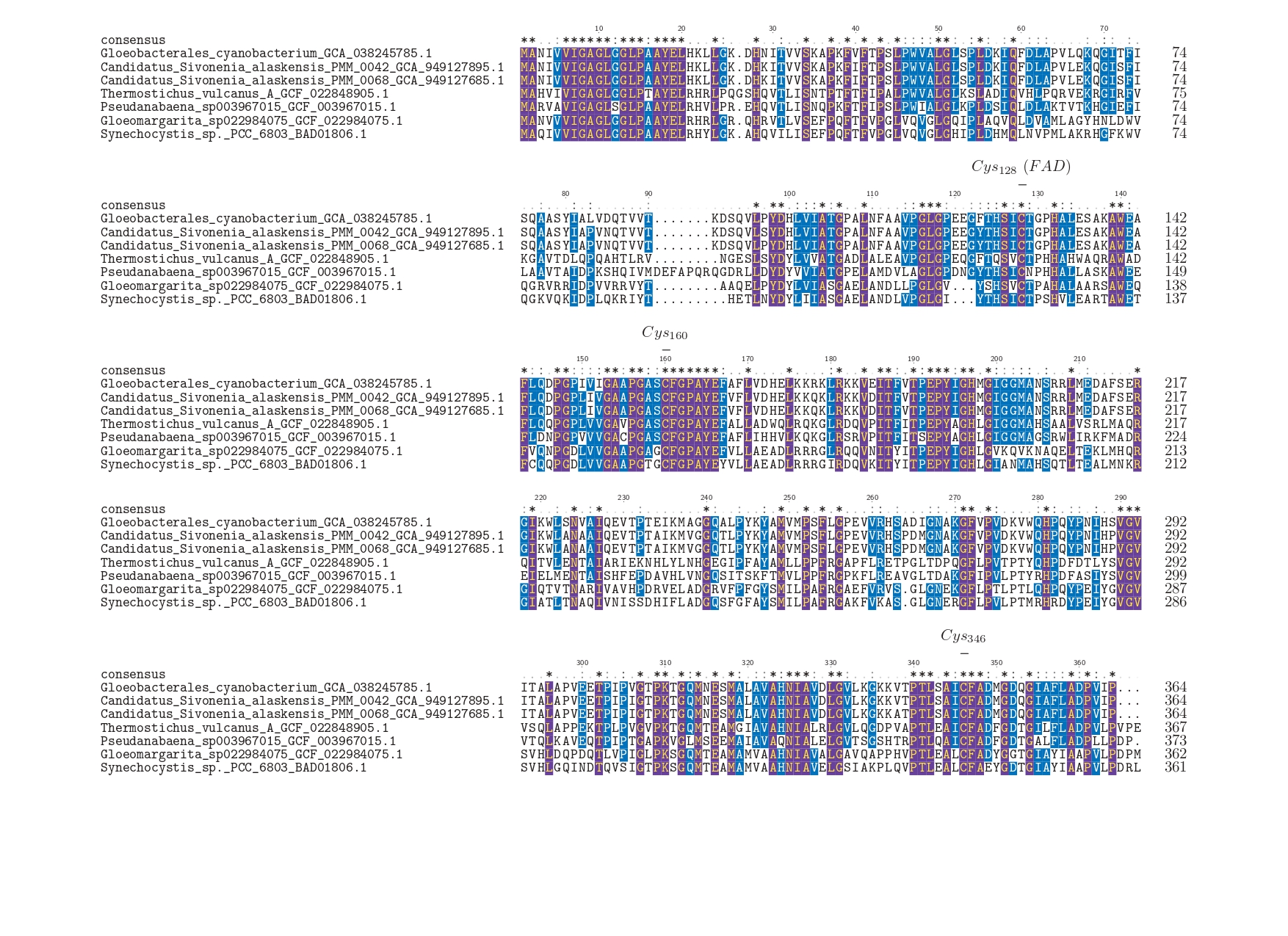

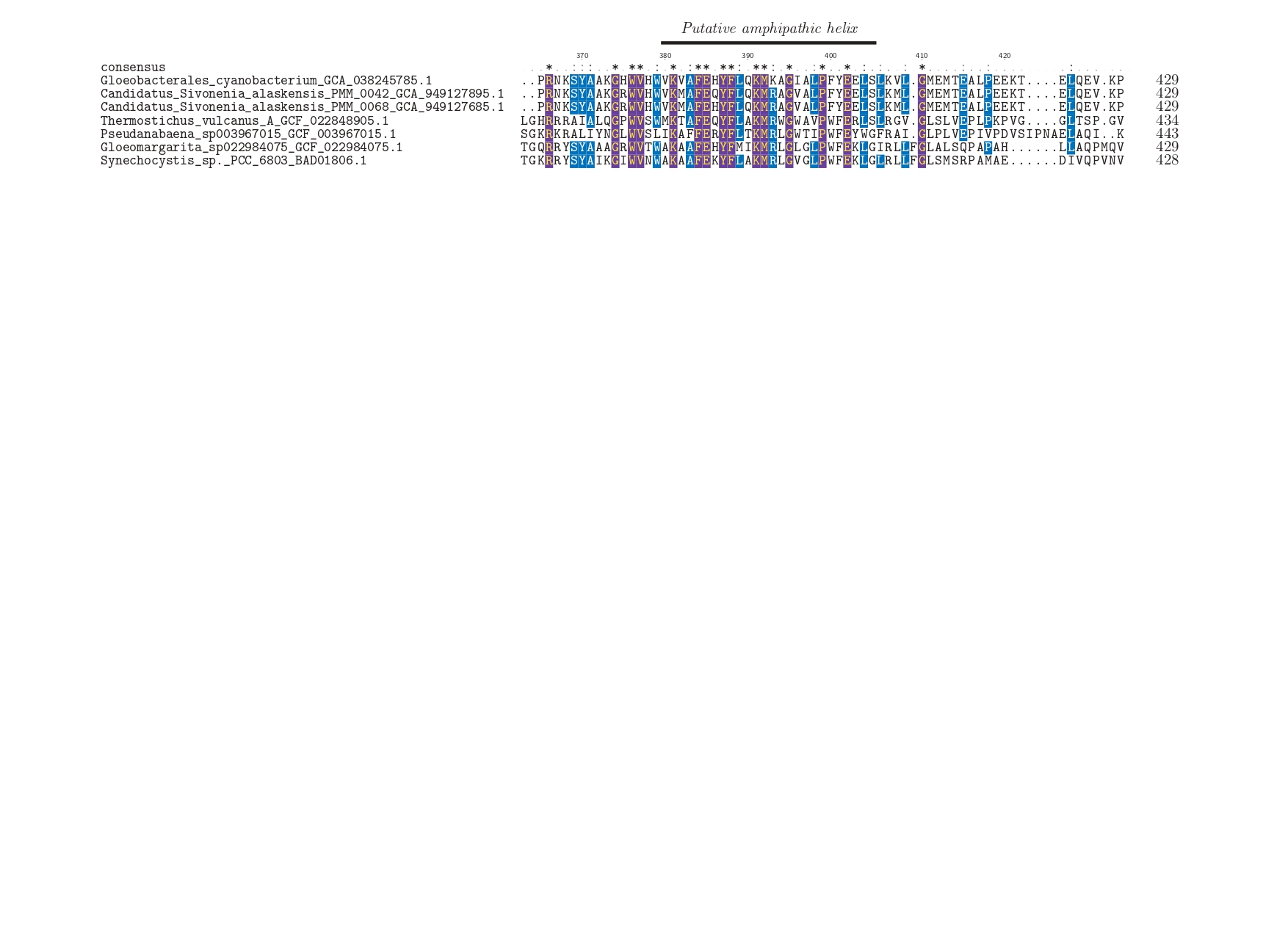

**Supplemental Figure 4: Amino acid sequence alignment of the three *Gloeobacter* SQR type I with SQR type I sequences of other basal lineages and the sequence of Synechocystis PPC 6803 substr. Kazusa Sll5036.** The residues playing a direct role in the catalytic mechanism of the enzyme, by covalently binding FAD or creating a sulfide bond, are annotated. The C-ter putative amphipathic helix supposedly anchors the protein within a membrane (10.1021/bi9003827). Residues that are 100% conserved are indicated by an asterisk (*); residues conserved in more than 50% of the sequences are indicated by a colon (:); and poorly conserved residues are indicated by a dot (·).

| **Accession** | **Biosample** | **Longitude** | **Latitude** | **Country** | **Cold criterion** |
| --- | --- | --- | --- | --- | --- |
| SRR7514503 | SAMN09633317 | -111.338851 | 58.778056 | Alberta, Canada | Subarctic (50°–66°N) |
| ERR10836081 | SAMEA112464045 | 158.8320278 | -54.71655556 | Macquarie Island, Australia | Subantarctic island (explicit polar region) |
| ERR10836083 | SAMEA112464047 | 37.58983333 | -46.94294444 | Marion Island, South-Africa | Subantarctic island (explicit polar region) |
| ERR12143029 | SAMEA114504931 | -73.41625 | 47.81176 | St. Maurice River, Canada | Temperate (<50°N, non-polar) |
| ERR12143032 | SAMEA114504934 | -73.41625 | 47.81176 | St. Maurice River, Canada | Temperate (<50°N, non-polar) |
| ERR3772646 | SAMEA6430891 | 102.289 | 72.399 | Tundra, Russia | Polar (>66°N, Arctic tundra) |
| SRR10595505 | SAMN13483817 |  |  | Artic | Polar (explicit Arctic location) |
| SRR3480285 | SAMN04958484 | -170.25 | 57.18 | Lake Hill Alaska | Subarctic (50°–66°N) |
| SRR3985421 | SAMN05421611 | -156.61 | 71.2999 | Barrow, Alaska | Polar (>66°N) |
| SRR3985424 | SAMN05422056 | -156.61 | 71.2999 | Barrow, Alaska | Polar (>66°N) |
| SRR3985527 | SAMN05422055 | -156.61 | 71.2999 | Barrow, Alaska | Polar (>66°N) |

**Supplemental Table 1 : Information on SRA identified by Logan search.**

| **Criteria** | **Cyanobacteria sampling** | **GTDB representative species** |
| --- | --- | --- |
| Maximum copy number | 3 | 10 |
| Maximum sequence coverage | 1 | 1 |
| Maximum –log(e-value) | 241 | 233 |
| Maximum sequence length | 515 | 646 |
| Minimum copy number | 1 | 1 |
| Minimum sequence coverage | 0 | 0 |
| Minimum –log(e-value) | 154 | 18 |
| Minimum sequence length | 282 | 215 |

**Supplemental Table 2 : Criteria applied for the selection of similar SQRI sequences using the Ompa-Pa tool.**

| genome | n_genes_called | n_genes_mapped | n_contigs | taxonomic_level | proportion_genes_retained_in_major_clades | genes_retained_index | clade_separation_score | contamination_portion | n_effective_surplus_clades | mean_hit_identity | reference_representation_score | pass.GUNC |
| --- | --- | --- | --- | --- | --- | --- | --- | --- | --- | --- | --- | --- |
| GCA_038245785.1 | 5142 | 4612 | 224 | phylum | 0.92 | 0.82 | 0.26 | 0.12 | 0.26 | 0.62 | 0.51 | TRUE |
| GCA_949127685.1 | 2942 | 2658 | 344 | phylum | 0.94 | 0.85 | 0.16 | 0.06 | 0.12 | 0.62 | 0.53 | TRUE |
| GCA_949127895.1 | 3042 | 2742 | 380 | phylum | 0.94 | 0.84 | 0.34 | 0.07 | 0.15 | 0.62 | 0.52 | TRUE |

**Supplemental Table 3 : GUNC metrics for contamination assessment of the three *Gloeobacter* MAGs.**

| Name | Completeness | Contamination | Completeness_Model_Used | Translation_Table_Used |
| --- | --- | --- | --- | --- |
| GCA_038245785.1 | 98.24 | 1.42 | Gradient Boost (General Model) | 11 |
| GCA_949127685.1 | 94.6 | 0.45 | Gradient Boost (General Model) | 11 |
| GCA_949127895.1 | 92.74 | 1.93 | Gradient Boost (General Model) | 11 |

**Supplemental Table 4 : Checkm2 metrics for contamination assessment for the three *Gloeobacter* MAGs.**
